# It matters who you are: Biography modulates the neural dynamics of facial identity representation

**DOI:** 10.1101/2025.04.08.647791

**Authors:** Yang Shi(施扬), Sophia Gommlich, Gyula Kovács

**Affiliations:** Department of Biological Psychology and Cognitive Neurosciences, Institute of Psychology, Friedrich Schiller University Jena, D-07743 Jena, Germany

**Keywords:** EEG, Familiarity, facial identity, Recognition, person knowledge, learning, multivariate pattern analysis

## Abstract

Recognizing the face of a person is an essential capacity to our social life. However, to interact properly with others we also need to know to which person a face belongs. Here we tested how person knowledge modulates face processing. Participants were familiarized with highly variable faces, associated with artificial biographies. Crucially, participants were allocated randomly into two groups, trained on the identical faces of the same persons, but with reversed person knowledge associated with the faces. Multivariate pattern analyses were used to examine the time course of identity representations in the EEG data. We estimated cross-participant, leave-one-participant-out pairwise identity classifications within the same face-person association groups and compared them to the those performed across association groups. We observed that face-person knowledge associations led to a robust familiarity signal from 300 ms and to a rapidly emerging identity representation, starting from 80 ms. Importantly, the shared associations within participant groups led to a longer-lasting and stronger face identity representation over the right posterior electrode cluster when compared to cross-group comparisons with reversed associations. The direct comparison of within and cross-group classifications showed that an early stage of identity representation, between 250 and 370 ms, is significantly modulated by face-biography associations. Our findings suggest that top-down, person recognition memory related information modulates visual face identity representation already at an early processing stage. Our study provides new insights into the spatiotemporal dynamics of how person-related conceptual, biographical knowledge modulates familiar identity representation.

## Introduction

Familiarization with previously unseen persons and recognizing them is an essential ability in social interactions. Prior studies showed that the processing of familiar faces has strong advantages over unfamiliar ones (Ambrus et al., 2021; Bruce & Young, 1986; Jenkins et al., 2011; Karimi-Rouzabahani et al., 2021; Visconti di Oleggio Castello & Gobbini, 2015; Young & Burton, 2017; for a review see Burton et al, 2011).

Several event-related potential (ERP) studies have investigated the temporal dynamics and cortical distribution of the familiarity effects and identified several components that are sensitive to face familiarity including the N250 and the “sustained familiarity effect” (SFE) manifest in the 400 - 600 ms time-window (for a review see Wiese et al, 2024). Later, multivariate pattern analysis (MVPA) studies revealed that familiarity information is indeed decodable in a time window, corresponding to these components (Ambrus et al, 2019, 2021; Dobs et al., 2019; Karimi-Rouzabahani et al., 2021; Li et al. 2022).

Importantly, familiarity cannot solely be attributed to perceptual exposure but can also be derived from additional non-visual contexts or person-specific knowledge, such as a name, occupation or personality traits (Eick et al, 2020; Schwartz & Yovel, 2016; Tsantani et al, 2021). When associated with newly familiarized faces, biographical information enhances face recognition and memory (Brooks et al, 2019; Mattarozzi et al., 2018; Schwartz & Yovel, 2016, 2019a, 2019b; Stolier & Freeman, 2016). This suggests that person knowledge enhances face/person recognition performance and modulates familiar representations through top-down mechanisms, extending beyond perceptual processing (Brooks & Freeman, 2019; Oh et al, 2021). However, the neurocognitive mechanisms underlying this modulatory effect are still not well-understood.

Neuroimaging studies have identified several brain regions sensitive to person knowledge, including frontal, medial and inferior temporal regions, as well as the amygdala and hippocampus (Bein et al., 2020; Borghesani et al., 2019; Brooks et al., 2019; Brosch et al., 2013; Cao et al., 2020; Cloutier et al., 2011; Collins & Olson, 2014; Gainotti, 2007; Goesaert & Op de Beeck, 2013; Stolier & Freeman, 2016; Tsantani et al., 2021; but see Verosky et al., 2013). The few available electrophysiological studies are inconclusive and showed semantic information sensitivity from 180 ms up to 600 ms, in several processing stages (Abdel Rahman, 2011; Gamond et al., 2017; Taylor et al., 2016; Wiese & Schweinberger, 2011, 2015). Thus, the temporal dynamics of the person knowledge modulation on facial identity (ID) remains relatively unclear as of today.

Therefore, we applied time-resolved cross-participant MVPA methods to the EEG data (Dalski et al., 2022; Grootswagers et al., 2017). We manipulated the artificial biography associated with newly learned faces during a training. Crucially, we allocated participants into two groups randomly, trained on the identical faces of the same IDs but with reversed associated person knowledge. For example, if the faces of one ID were associated to the biography of “Anna” and the faces of another ID were associated to “Louise” in one group, the face - biography associations were reversed for the participants of the other group (the faces of the first ID were associated to the biography of “Louise” while the second to “Anna”). In a subsequent EEG session, we recorded the brain responses and analyzed the cross-participant pairwise decoding accuracy of the familiarized IDs from within the same face-person association group and compared it to the decoding across the two association groups. Thus, we compared face representations sharing perceptual and person specific knowledge with representations having different associated person knowledge. We found that face representations are sensitive to the associated person knowledge and the shared associations within participant groups increased decoding performance between 250 and 370 ms when compared to the reversed associations in the cross-group comparisons. This approach allows us to study the temporal dynamics of the modulation of person knowledge on face representation and opens a novel way of testing the differences of face and person representations.

## Materials and Methods

### Participants

A total of 45 participants (7 males) with an average age of 22.4 years (SD = 3.64) participated in the experiment in exchange for partial course credits. They were allocated randomly into two groups: *Group 1* (n = 22) and *Group 2* (n = 23). Participants reported no history of neurological conditions, had normal or corrected-to-normal vision, and were all right-handed except three individuals. The experiment was conducted in accordance with the guidelines of the Declaration of Helsinki and was approved by the ethics committee of the University of Jena. Written informed consents were acquired from all participants.

### Stimuli

The face stimuli consisted of 20 highly variable, colored ambient images of eight IDs (four males) each. All IDs were foreign celebrities whom participants were unfamiliar with. Four IDs were selected for the familiarization learning (two males, two females; see *below*). Stimuli were cropped and resized to 400 × 400 pixels, and placed on a unified neutral-grey background. Stimuli were presented on a LCD display with a resolution of 1920 × 1080 pixels and with a refresh rate of 120 Hz. The experimental programs were written in Psychopy (Peirce et al., 2019).

As person associated knowledge, four biographical information sets were created. These biographies were fully fictional and did not relate to any real individuals. Each biography described personal information about a fictional individual either from France or Germany, including their name, age, nationality, occupation, hobbies and personality traits (for examples see Figure 1). The full text of the training biographies can be found in Supplementary Materials (Appendix 1). During training, each biography was associated to the faces of one of the four IDs for each participant in the following way.

**Figure 1.**
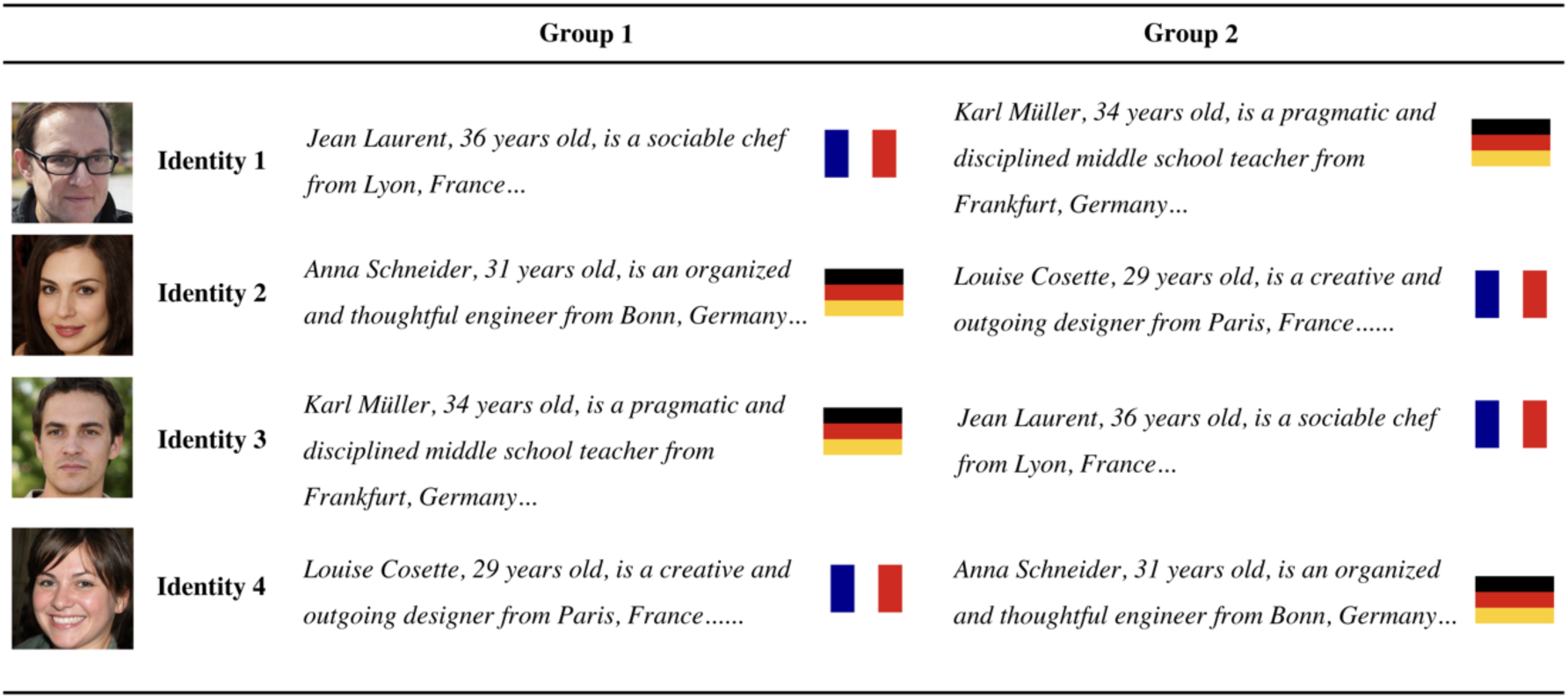
The manipulation of face and semantic person knowledge association for the four trained identities and two participant groups, separately. A sample face image of each identity (ID) is presented on the left. Next, we present a sample of the artificial biographies of the training phase, together with national flags, signaling the country of origin of the persons, for better understanding, separately for the two participant groups. For the full version of person biographies see the Supplementary Materials (Appendix 1). Please note that participants were allocated randomly to either group and that the presented faces were identical for every participant, but they were associated with reversed biographical information. This manipulation of face-person knowledge association enabled us to keep the visual perceptual stimulation constant, while reversing the associated person knowledge across participant groups. Example faces were created by AI for illustration purposes (https://thispersondoesnotexist.com/).

First, we allocated participants randomly into two face-person association groups. Importantly, the association between the faces of the IDs and the associated person knowledge was reversed across the two groups. Figure 1. illustrates the face-person knowledge associations for the two participant groups. For example the faces of ID1 were associated to the artificial biography of “Jean, a French male” while the faces of ID3 were associated to “Karl, a German male” in one group while the face - biography associations were reversed for the participants of the other group (i.e. the faces of ID1 were associated to the knowledge about “Karl” and the faces of ID3 were associated to “Jean”). Participants were not aware of the two possible association groups.

The familiarization training of Group1 and Group 2 was identical, except for reversed face-person knowledge association.

### Procedure

Participants completed four sessions (three training and a final EEG recording session) over three consecutive days. On the first two days, participants completed a 15-minute training session each day. On the last day, they completed a third training session, immediately followed by the EEG recording.

Each training session included two parts: a learning and a testing phase. During the learning phases participants were presented five different face images of each of the four training IDs. Each training session introduced five new face images per ID. The faces were presented above the associated biographical information. The background biographical information was progressively revealed as participants navigated through five image + biography screens (Figure 2). Participants could freely navigate through the screens using a keyboard without any time limit and were instructed to memorize the faces and their biographies as good as possible.

**Figure 2.**
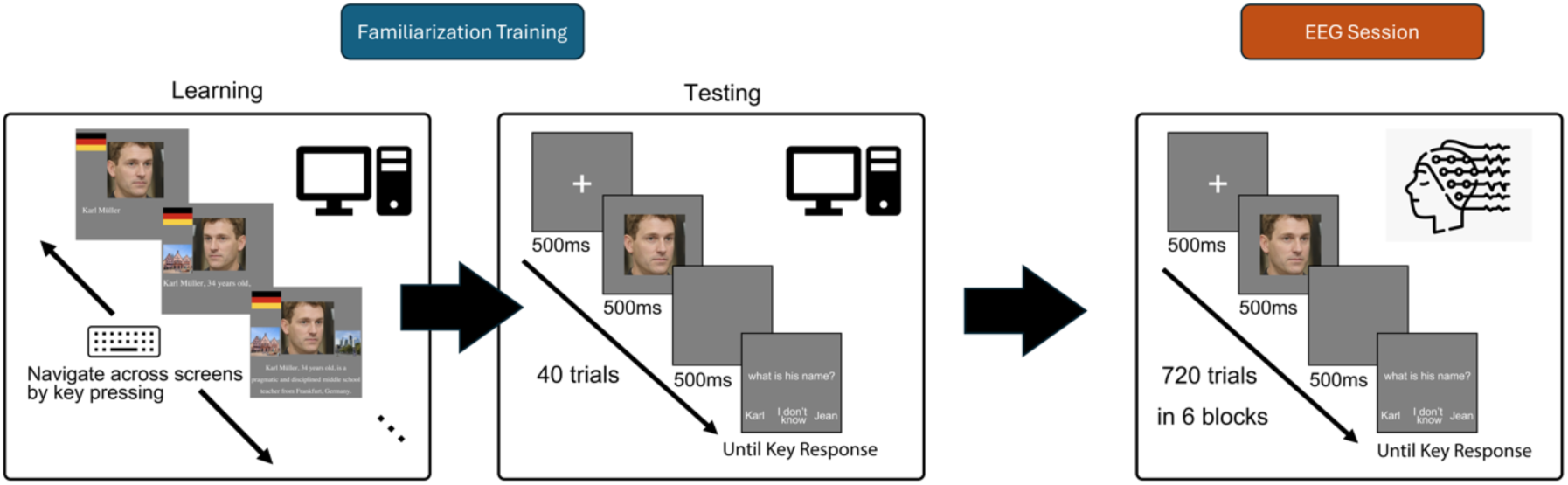
The Experimental paradigm. The experiment was composed of a three-day long familiarization training and a post-familiarization EEG recording session. The familiarization training was performed on three consecutive days and was composed of two phases, a learning and a testing phase. The EEG was recorded during the testing phase of the third day, immediately following the last learning phase. For details see Methods section.

Once participants finished the learning phase, the second, testing phase commenced. During testing, participants had to complete 40 trials, which included 20 face images of the four learned identities, and 20 faces of four unfamiliar IDs. These unfamiliar faces served as control images as they enabled us estimating the face-biography associations better by measuring false alarms and correct rejections. Each testing-phase trial started with a 500 ms fixation cross, followed by a 500 ms blank and a face at the center of the screen for one second, followed by a 500 ms blank. Next, a question regarding the associated biography appeared in the middle of the screen. These questions could cover all five dimensions of the biographies: name, homeland, job, hobbies or personality traits (for example: *“What is his/her job?”* or *“Where does he/she come from*?”; for the complete list of questions please see Supplementary Materials Appendix S2). Three response options appeared below the questions: the two alternative answers (the correct one and the one from the other same-gender ID) at the left and right of the screen and the phrase “I don’ know” in the middle. Participants had to signal their answers by pressing a button and received a visual feedback: the correct alternative turned green for correct answers while the correct alternative turned green, and the incorrect choice turned red for incorrect answers.

Importantly, different biographical information was assigned to the faces of a given ID in the two participant groups. With this approach, we created familiar ID representations with the same visual information but with reversed person knowledge across the two participant groups (see Figure 1).

### EEG Recording and Preprocessing

The EEG recording session consisted of six blocks of 120 trials each (720 trials in total) and lasted approximately 50 minutes. During the EEG recording, participants performed the same paradigm as in the testing phase (Figure 2). Face images of eight IDs (four familiarized and four unfamiliar IDs) were presented. For each ID, five new, previously unseen images were introduced to prevent image-level learning effects. The EEG was recorded in a dimly lit, electrically shielded and sound-attenuated chamber. The distance between the eyes and the computer screen was 96 cm and participants’ head layed on a chin rest. The EEG was recorded using a 64-channel Biosemi Active II system, with a common mode sense (CMS) and drive right leg (DRL) ground reference (sampling rate: 512 Hz). The preprocessing pipeline was implemented in MNE-Python (Gramfort, 2013). EEG signals were visually inspected to identify bad channels and artifacts such as extreme amplitude fluctuations. Bad channels were interpolated using built-in functions of MNE-Python. The EEG data was re-referenced to the average of all electrodes, a 50 Hz notch filter was applied to remove power line noise, followed by a band-pass filtering between 0.1 Hz and 40 Hz, using a zero-phase finite impulse response (FIR) filter. An independent component analysis (ICA) algorithm was applied to the data to find and remove ocular artifacts (Iriarte et al., 2003), based on the topographic distribution and time course of the components. Finally, the data was segmented from -200 to 1300 ms relative to stimulus onset, and baseline corrected to the first 200 ms of the epochs. The resulting data was then down-sampled to 100 Hz to improve signal-to-noise ratio for the MVPA.

### Multivariate Analysis (Decoding)

#### Familiarity representation

To confirm if participants were indeed familiarized with our stimuli first a representational similarity analysis (RSA) was conducted on the EEG data (Figure 3a). For each participant, at each time point, neural representational dissimilarity models (RDMs) were constructed using neural dissimilarity measures, computed from all face image pairs (8 IDs, 5 images per ID, 40 images in total). Specifically, dissimilarity measures were calculated by performing pairwise decoding on all face image pairs using Support Vector Machine (SVM) classifiers with radial basis function (RBF) kernels (Scholkopf et al., 1997). Classifiers’ training and testing were performed across all electrodes of the scalp as well as independently for 6 predefined regions of interest (ROIs; Ambrus et al., 2019; 2021) using a 10-fold cross-validation approach: the classifiers were trained on 90% of the datasets and tested on the remaining 10% of the trials in each iteration. The procedure was repeated until each trial had been used as training and testing set once. The decoding accuracies were then averaged across iterations.

**Figure. 3.**
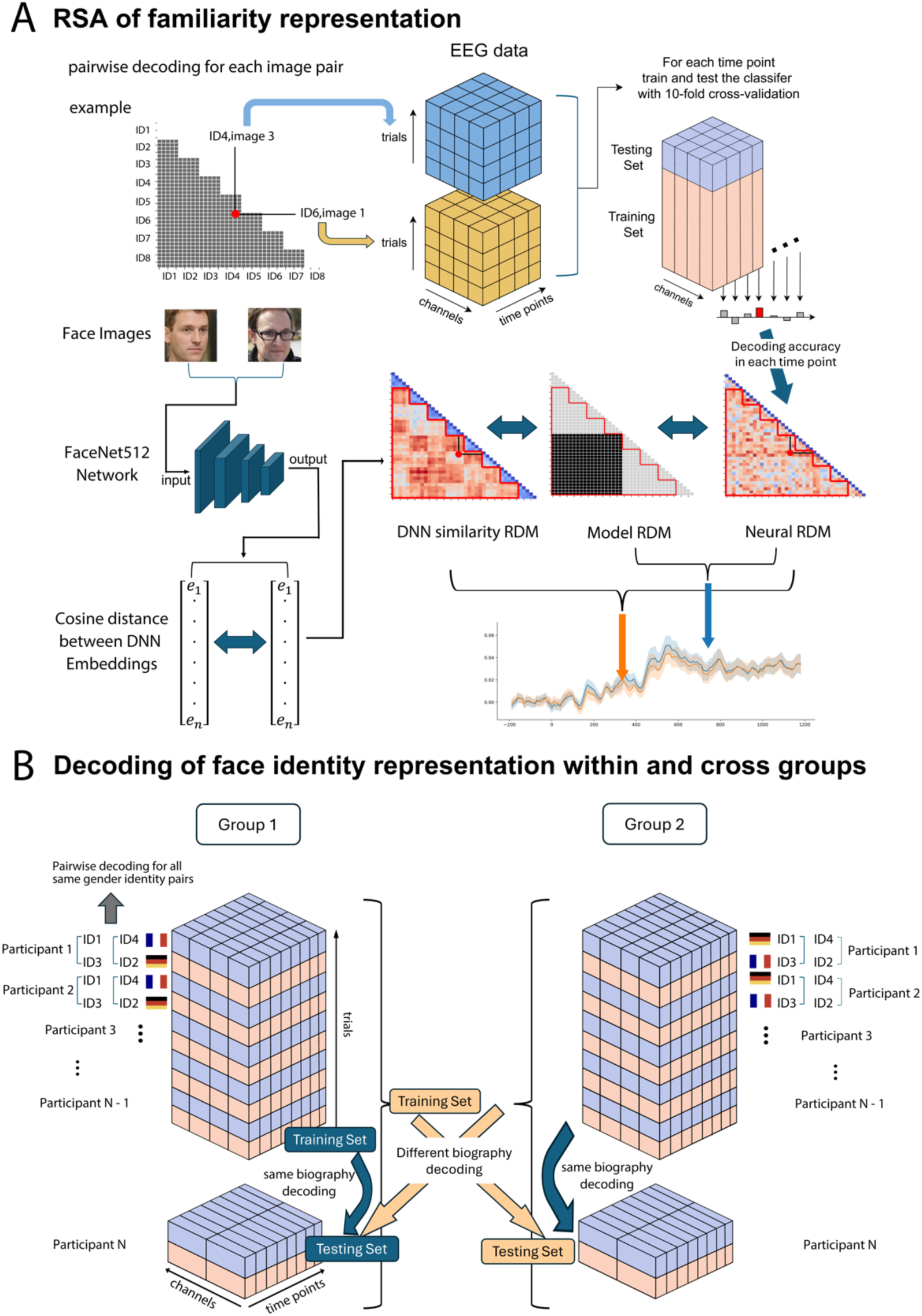
The logic of the multivariate pattern analysis. **A.** Familiarity decoding using RSA. SVM classification analyses were performed for each combination of individual images, using a 10-fold cross-validation scheme for each time point separately. This procedure led to a 40 × 40 matrix (i.e., 5 images for each of the 8 identities) of decoding accuracies at each time point. The resulting decoding accuracy values were used to construct a neural RDM for each time point (right). To assess the familiarity effect, we computed the correlation between the neural RDM (right) and the model RDM (middle) which encodes the predicted familiarity effect for each participant and time-point separately. To control the effects of low-level face similarity, we calculated the distance of DNN embeddings between each pair of faces and constructed the similarity RDM, using the distance information as well. We then performed a partial correlation analysis between neural and model RDMs while partialling out the DNN similarity RDM (left). Notably, the correlation analyses included only image pairs that did not belong to the same IDs (red frames of the RDMs) **B.** The comparison of identical (within group) and reversed (cross-group) face-person-knowledge associations. For within-group decoding, accuracy was computed using a leave-one-participant-out cross-validation approach from both groups separately, for each pair of same gender faces. For cross-group decoding, the classifier was trained on the participants of one group and tested on the participants of the other. We reasoned that within group decoding reflects both shared visual and biographical information, while the biographical information is not shared in the cross-group comparisons. For details see Methods section.

To model the familiarity effect, we created a predictor RDM with 40 × 40 cells. Each cell was assigned the value of zero for comparisons of similar images (both images are familiar or unfamiliar), while a value of one was assigned to comparisons of dissimilar images (one image is familiar while the other is unfamiliar). To quantify the correlation between predictor and empirical RDMs, we vectorized both RDMs by extracting the lower off-diagonal values and calculated the Pearson correlation coefficient between the two extracted vectors at each time point, electrode cluster and participant separately. This procedure led to a time series of correlations that quantifies the correspondence between neural representations and the predicted pattern (Kriegeskorte, 2008). To avoid confounding familiarity information with identity information, all comparisons within the same identity (i.e., two different images of the same ID) were excluded from the analysis (Ambrus, 2019).

In order to control for the potential effects of facial similarity, we used a deep neural network (DNN; Schroff et al., 2015) assuming that it captures the facial similarity beyond pixel-based similarity (Dobs et al., 2022). All face image pairs were processed through the DNN embedding pipeline, which includes face detection (with the RetinaFace algorithm), automatic alignment, and normalization. Next, the DNN embedding vector for each image was extracted from the final, fully connected layer of a pre-trained FaceNet-512 network, leading to a 512-dimensional feature representation. The DNN-similarity RDM was created by calculating the cosine distances between all pairs of embedding vectors. All steps, including face detection, alignment and feature extraction were implemented in DeepFace package (Serengil & Özpınar, 2024) using Python.

#### Face Identity representation within and across person-knowledge association groups

To estimate the modulatory effect of biographical information on face representations, we compared the decoding accuracy of same-gender familiar face ID pairs within as well as across person knowledge association groups (Figure 3b). First, we performed a time-resolved decoding analysis within each group (training and test datasets originating from the same group). Specifically, we applied SVM with RBF kernel to classify same-gender face ID pairs for participants of Group 1. Decoding accuracy was calculated by using a leave-one-participant-out (LOPO) cross-validation approach, by training the classifier on the data of N – 1 participant and testing it on the remaining participant of the same group in each iteration until each participant in has been tested once. The decoding accuracies were averaged across pairs and participants. The same analysis was conducted independently for the participants of Group 2 as well. Finally, these two within-group decoding accuracies were averaged.

Next, we conducted another LOPO decoding analysis across the two face-person association groups (training and test datasets originating from different participant groups). To construct the training set, we concatenated the corresponding data of the participants in one group for the same ID pairs as previously. The classifier was then trained on this dataset and tested on the participants of the other group, and iterated until every participant of the other group was tested once. The resulting time-resolved decoding accuracy was then averaged across ID pairs and participants. The same analysis was then repeated in the opposite direction (training and testing groups were exchanged). Finally, the cross-group decoding accuracy was calculated by averaging the accuracies from both training-testing directions (Group 1 → Group 2 and Group 2 → Group 1).

To further investigate the stability of neural ID representations over time, we performed a temporal generalization analysis within as well as across person-knowledge association groups. This analysis was identical to the above described one, with the exception that the classifier was trained at each time point of a trial in the training set and tested on all the time points of a trial in the testing set, resulting in a two-dimensional temporal generalization matrix. In the matrix, a significant decoding along the diagonal suggests transient, sequential processing stages, while a broader, temporally sustained off-diagonal decoding indicates temporal generalization and a stable neural code shared between training and testing sets over time (King & Dehaene, 2014).

### Statistical testing

To increase the signal-to-noise ratio, all analyses were conducted on data down-sampled to 100 Hz, and a moving average of 30 ms (3 consecutive time-points) was applied to all decoding accuracy data at the participant level (Ambrus et al., 2019; Dalski et al., 2021; Kaiser et al., 2016). Two-sided cluster-based permutation tests with 10,000 iterations were conducted on all the measurements to identify significant effects. Clusters having *p* < 0.01 were marked as significant. The decoding and statistical analysis were conducted using Python, MNE-Python, scikit-learn and SciPy package (Virtanen et al., 2020).

## Results

### Behavior

Behavioral performance was analyzed by calculating accuracy and response times (RT) of the EEG recording session. Participants demonstrated high-level of accuracy in the face-biography matching task (M = 0.98, SD = 0.01). Next, we calculated *d’* in the following way. A “hit” was defined as a correct response in familiar ID trials, and a “correct rejection” was defined as the “I don’t know” response in unfamiliar ID trials. Then “false alarm” rates were calculated by 1 – correct rejection rate. Then, the hit and false alarm rates were z-transformed to Z (hit) and Z (false alarm). Finally, the *d’* was calculated according to the equation of Stanislaw and Todorov (1999):

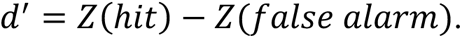

One sample t-test revealed that participants’ sensitivity was significantly larger than zero (M = 5.34, SD = 1.23; *t*_(37)_ = 26.88, *p* < 0.001, Cohen’s d = 4.36). Finally, response times were significantly longer for familiar (M = 1.74 s, SD = 0.31) when compared to unfamiliar IDs (M = 0.98 s, SD = 0.17; paired t-test: *t*_(37)_ = 20.562, *p* < 0.001, Cohen’s d = 3.34. Altogether, these results suggest that participants were well familiarized with the faces and learned their association to the person related knowledge well.

### Familiarity representation

To uncover the cortical distribution of familiarity representation of the learned faces across time, we performed an RSA, both on the EEG data of the entire scalp as well as in six pre-defined topographic ROIs, including the right and left frontal, central-temporal and parietal-occipital cortex (Figure 4, blue lines).

**Figure. 4.**
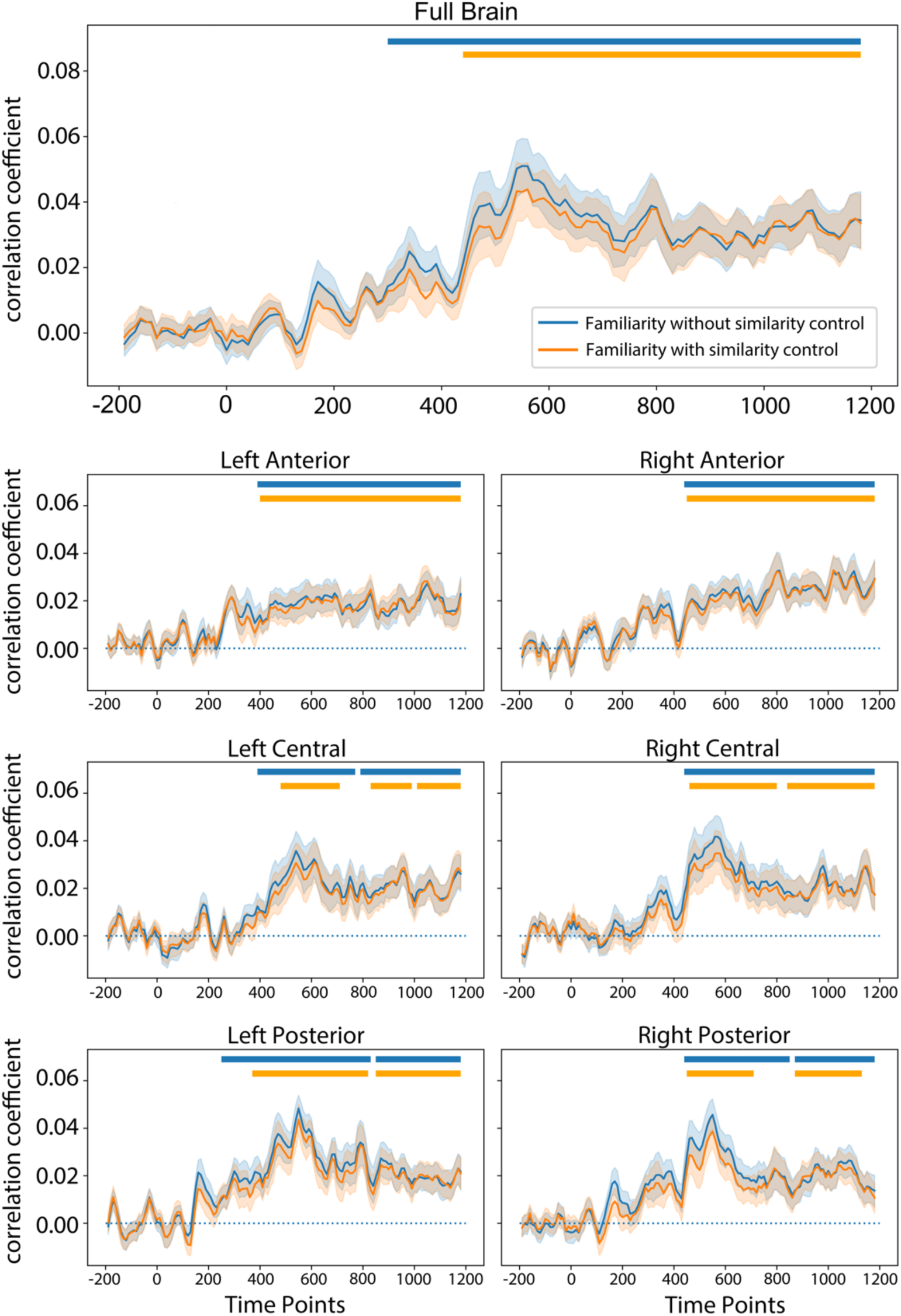
Neural representations of face familiarity. Time-resolved correlations between neural and familiarity predictor RDMs for the entire scalp (top) and for each of the six ROIs separately. Blue lines represent correlations without partialling DNN-based similarities out, while yellow lines represent correlations with partialled out DNN-based image similarities. Horizontal lines indicate statistically significant time windows (p < 0.01 with two-tailed cluster-based permutation test against zero correlation). Shaded regions represent ± 1 SEM.

When testing the response patterns across the entire scalp, significant correlations emerged between the familiarity model RDM and the neural RDMs from 300 ms, peaking at 550 ms and remained significant until 1200 ms. RSAs in the pre-defined ROIs revealed significant familiarity effects very similarly in each electrode cluster (for details see Supplementary Materials, Table 1) These results strongly suggest that our familiarization procedure indeed effectively induced a different neural representation for the faces of the familiarized IDs, across the entire scalp.

To ensure that the observed familiarity effects are not confounded by low-level factors, we measured visual similarity between face image pairs using deep neural network (DNN) embeddings and partialed out their effects from the familiarity decoding results.

After controlling for face similarity, the significant familiarity effects appeared later (onsets from 440 ms, peak: 560 ms) over all electrodes as well as over left central and posterior ROIs (Supplementary Materials, Table 2). These results indicate that the earliest familiarity representations, emerging from around 300 ms to 440 ms are affected by low-level face similarity, as controlling for DNN similarity reduced the effect time window. In contrast, the familiarity representations after 440 ms are robust and not affected by similarities, captured by DNNs. Together, these results suggest that familiarity indeed developed during the three-day training period and that the later representations remain robust when controlling for low-level visual face features, reflecting familiarity, per se.

### Face Identity representation within and across person-knowledge association groups

For the first time, in our paradigm we associated the faces of given individuals to either the same (within association groups) or reversed (across association groups) person specific knowledge. This manipulation enabled us keeping the visual stimuli constant while testing the modulatory effect of altered biographic person knowledge. We hypothesized that if a representation is sensitive to the face associated person knowledge, then the shared associations within participant groups might increase the strength of leave-one-participant-out (LOPO) decoding performance when compared to the reversed associations in the cross-group comparisons. Alternatively, if face-specific ID representations are encoded independently of biographical person knowledge, then within and cross-group decoding performances should be similar, as the visual face information is identical in the two groups.

Figure 5 presents the results of time-resolved within- and cross-group decoding analyses for the four familiarized IDs. As a reminder, within-group LOPO classification results show the neural dynamics of ID representations when the semantic descriptions are identical across participants. These within-group decoding accuracies (Figure 5 blue lines) were strongly and robustly significant from 100 – 430 ms and 490 – 810 ms, (peak: 130 and 500 ms) when tested across the entire scalp.

**Figure. 5.**
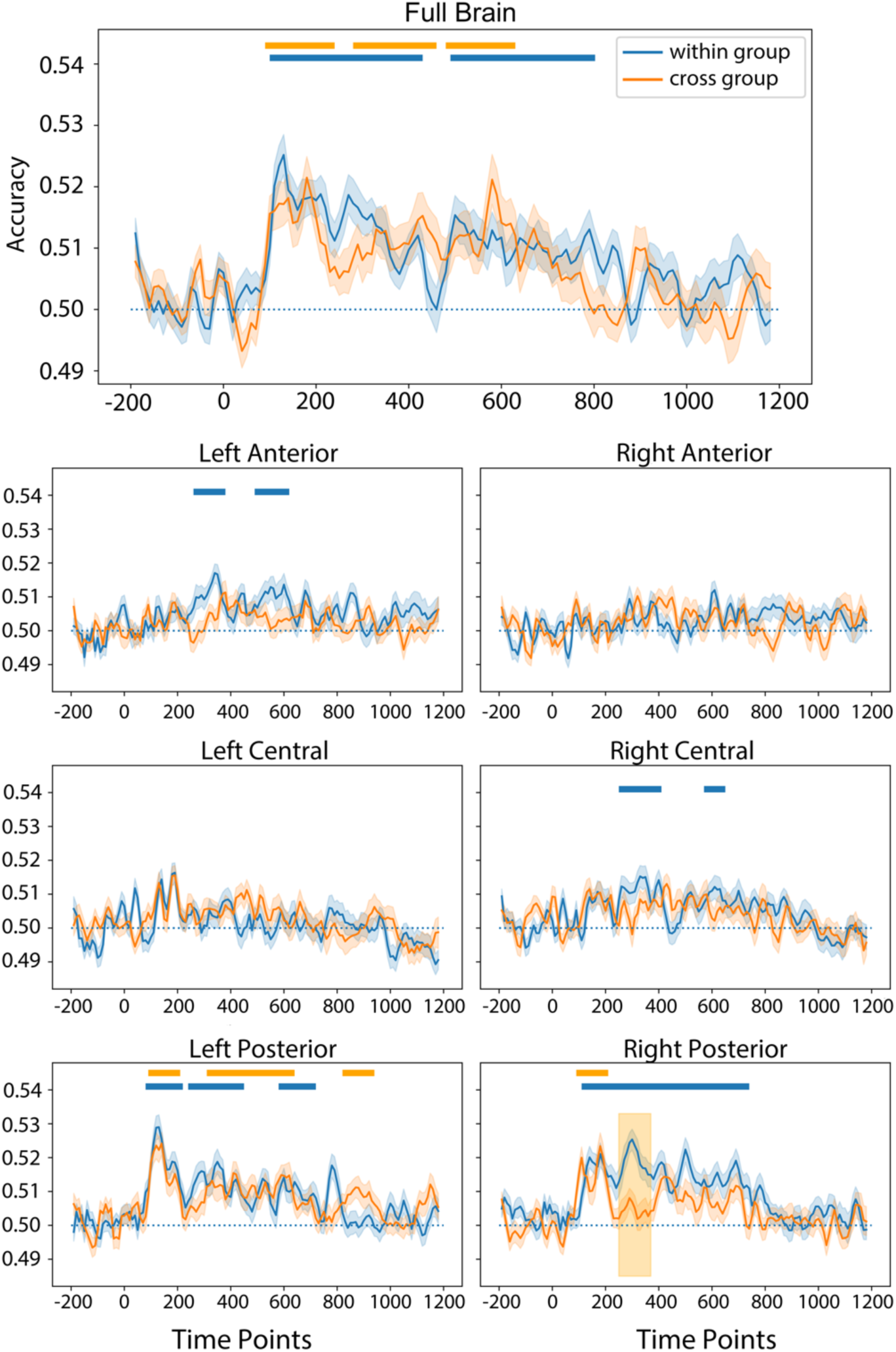
Time-resolved leave-one-participant-out face ID classification with shared and reversed biographical information separately. Biographical information is shared across participants for the within-group decoding (blue line) while it is reversed with respect to face IDs in the cross-group decoding (yellow line). The LOPO decoding performance for same-gender ID pairs is shown as the function of time for the entire scalp (top) and each of the six ROIs separately. Horizontal lines indicate statistical significance against chance level (p < .01, two-tailed cluster-based one-sample permutation test). The orange-shaded square region represents the statistically significant difference between the two curves (p < .01, two-tailed cluster-based permutation test). Shaded regions around the curves represent ± 1 SEM.

The earliest and strongest ID representation was found over posterior electrode clusters, where it emerged at 80 ms (left) and 110 ms (right) and remained significant until 720 ms (left) and 740 ms (right), with brief interruptions between 220-240 and 450-580 ms over the left hemisphere (peaks: 130, 350 and 600 ms for the left, and 300 ms for the right hemisphere, respectively). This suggests that ID specific representations are localized bilaterally over the occipital-temporal cortex.

Cross-group ID classification (Figure 5 yellow line) shows where and when in the brain the visual ID information is represented. As in this case person specific background information is different across participants, a significant decoding originates most likely from perceptual processing steps. Over all electrodes, significant LOPO cross-group decoding accuracies were observed somewhat weaker than for the within-group classification (onset: 90 ms and remains significant shorter, until 630 ms with interruptions between 240-280 and 460-480 ms; peaks: 180, 430 and 580 ms). In frontal and central regions, no significant decoding accuracies were observed in this case. In posterior electrodes, cross-group LOPO decoding accuracies remained significant, but the durations were strongly reduced, when compared to within group classification results: Significant time windows are found from 90 – 210 ms, 310 – 640 ms and 820 – 940 ms over the left, and only between 90 -210 ms over the right hemisphere (peaks: 140, 410 and 870 ms for the left, and 180 ms for the right hemispheres).

Most importantly to the conclusions of the current study, when comparing LOPO decoding accuracies within- and across person knowledge association groups, we found significantly stronger decoding in the time window between 250 - 370 ms over the right posterior electrode cluster (10,000 iterations two-tailed cluster-based permutations, *p*< .01)

This result confirmed our initial hypothesis and suggests that the shared associations within a given participant group enhance the strength of available ID information when compared to the reversed associations in the cross-group comparisons. Therefore, one can assume that the representation within the relatively early time-window of 250 and 370 ms post stimulus onset, is indeed sensitive to the face associated person knowledge and this effect is localized to the occipito-temporal electrode sites of the right hemisphere.

### Temporal generalization of ID neural representation within and across person-knowledge association groups

To investigate the temporal dynamics and stability of face ID representations within and across person-knowledge groups, we conducted a temporal-generalization analysis using the same group-level decoding procedure as the previous decoding analysis (see *Methods*). Figure 6 presents the generalization matrices for both within- and cross-group analysis. The within-group generalization on full-scalp data revealed two distinct stages during the unfolding of ID representations. During the first stage (100 – 160 ms), classifiers showed significant decoding accuracy along the diagonal, with only minimal off-diagonal generalization, suggesting rapidly evolving, time-specific ID neural representations. In the second stage (160 – 800 ms), classifier performance generalized across a broad range of time with a few interruptions, indicating a transition to more stable and possibly higher-level ID representations.

**Figure 6.**
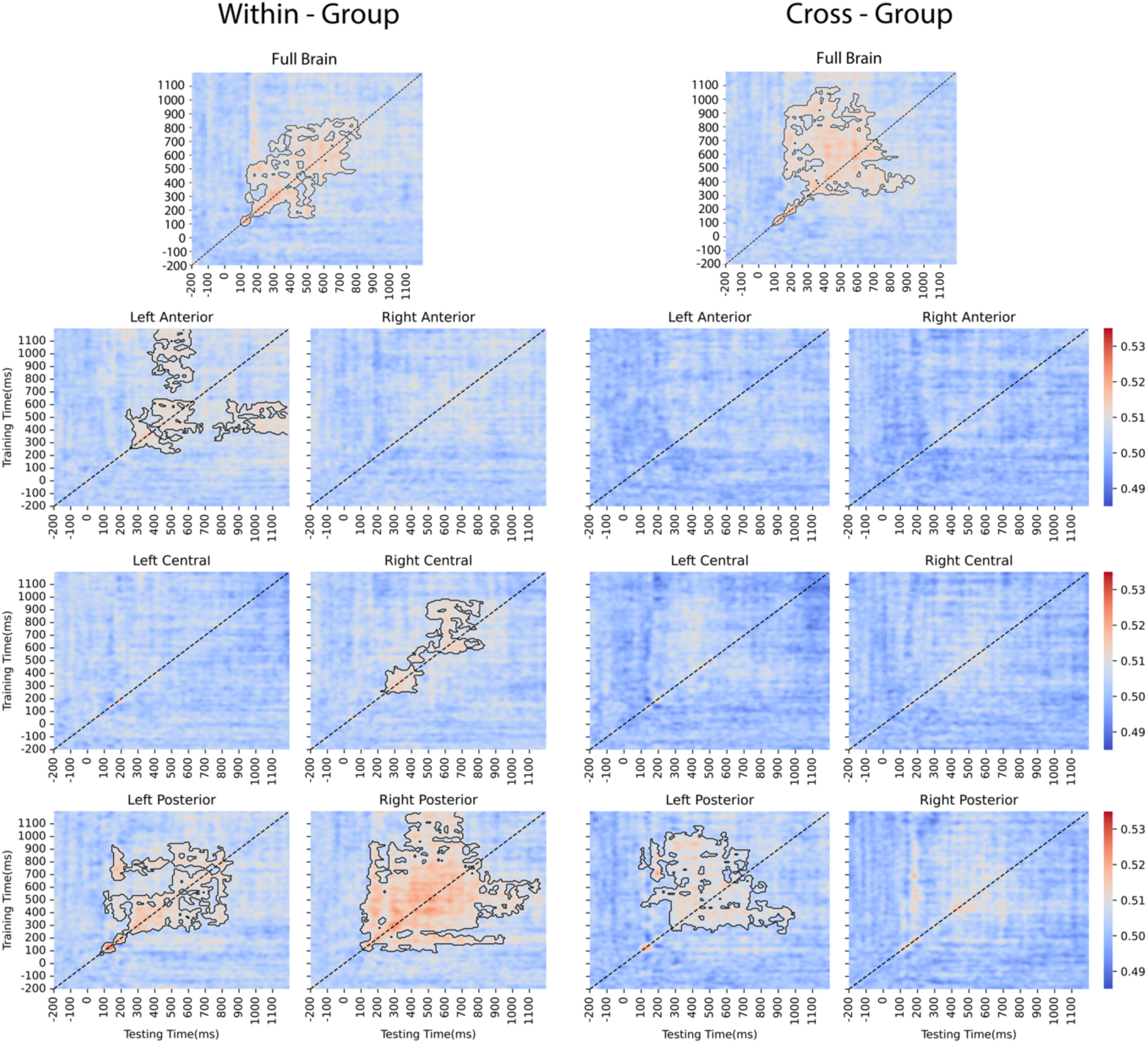
Temporal generalization of leave-one-participant-out face ID classification with shared and reversed biographical information, separately. Biographical information is shared across participants for the within-group temporal generalization analysis (left) while it is reversed with respect to face IDs in the cross-group analysis (right). Each classifier was trained on one time point of the training set and tested on all time points of the testing set (see Methods) for the entire scalp (top), and for each of the six ROIs, separately. Decoding accuracy is indicated by the color scale. Statistically significant spatial-temporal clusters are marked by black outlines against chance level (p < .01, two-tailed cluster-based one-sample permutation test). Black dotted lines indicate the diagonals.

The temporal generalization analysis revealed distinct patterns across the ROIs. For within-group comparisons, in bilateral posterior regions, classifier performance showed strong off-diagonal temporal generalization; however, the effect emerged with different latencies (from 250 – 800 ms in the left and 110 – 810 ms in the right hemisphere). Additionally, an early significant diagonal decoding from 80 – 250 ms with minimal off-diagonal generalization was observed in the left posterior region but not in the right. These results suggest that posterior regions have stable ID representations bilaterally, but they may differ in the early-stage ID processing. The right central ROI showed a different temporal generalization pattern from this. Specifically, significant diagonal decoding was observed from 250 – 420 ms and from 570 – 670 ms, with limited temporal generalization, suggesting transient changes of ID representations. Finally, in the left anterior region, diagonal decoding was observed similarly, from 260 – 380 ms with limited off-diagonal generalization and it was followed by a later diagonal decoding from 430 – 620 ms which generalized well over time (from around 750 ms to the end of the trial). This pattern suggests a sustained ID representation in the left anterior ROI during later processing stages.

The temporal generalization analysis across association groups revealed a two-stage temporal dynamics of ID representations over the full scalp as well. A significant diagonal decoding was observed from 90 – 300 ms, showing minimal off-diagonal spread, whereas the later diagonal decoding (from 300 – 700 ms) exhibited strong generalization over time. Compared to the pattern of within-group generalization on the full scalp data, cross-group temporal generalization was reduced between 160 – 300 ms, suggesting a disruption of cross-group ID representation stability, which is probably caused by the modulation of face-person associations.

For cross-association group analysis, only the left posterior ROI showed significant diagonal decoding from 390 – 670 ms. This showed a strong temporal generalization and occurred within a similar time window as in the within-group analysis. These results suggest that ID representations in the left posterior region may be less affected by person-related semantic knowledge when compared to the entire scalp or other ROIs.

## Discussion

We used EEG and MVPA with leave-one-participant-out cross-validation to study the modulation of the spatial-temporal dynamics of face ID representations by person knowledge. The major findings of the study are the followings. (1) A brief face – biography association training led to long-lasting familiarity representation from ∼ 300 ms post-stimulus onset. (2) Familiarity representation prior to ∼ 400 ms was affected by face similarity. (3) Face ID information emerges around 80 ms over the right posterior electrodes and when tested for participants with matching face - biography information remains significant long. (4) Most important to the aims of our study, when estimated across participant groups, having reversed face-biography associations, ID information was significantly shortened and its direct comparison with and without shared biographies showed that an early processing stage (250 - 370 ms) in right posterior region is modulated by person specific biographical knowledge significantly. This result is further confirmed by the temporal generalization of the results, which indicated reduced cross-group decoding accuracies.

### Familiarity representation after face-biography association training

Familiarity representation was manifest in a prolonged period from 300 ms onwards, across the entire scalp. This suggests that familiarity representation could emerge as the result of a brief face-biography association training, in line with previous studies, applying three-day long personal familiarization (Ambrus et al, 2021; Sliwinska et al., 2022). The relative late onset of familiarity information is in accordance with the results of previous studies (Ambrus et al, 2021; Wiese et al., 2019a, 2019b) as well.

Compared to previous magneto- and electrophysiological MVPA studies, we observed significantly broader distribution of familiarity representations. For example, a three-day long personal familiarization led to familiarity representations in a similar time-window (400 to 1000 ms), but restricted to bilateral posterior electrode sites (Ambrus et al., 2021).

It is plausible that the broader distribution in our current study is attributed to the explicit training of face-person knowledge associations, leading to stronger familiarity representations than the simple, unscripted personal meetings of prior studies. This refines previous conclusions that emphasized the role of longer, real-life personal familiarization methods (Ambrus et al., 2021; Sliwinska et al., 2022; Wiese et al., 2019a; Popova & Wiese, 2023a, 2023b; for a review see Wiese et al., 2024). Instead, the observed strong familiarity representation suggests the dominant role of person knowledge in obtaining familiarity with a person, independently whether familiarization happens digitally or in real-life. Specifically targeted experiments, comparing these familiarization methods are required to confirm this conclusion.

Further, the broad distribution of familiarity representation is consistent with multivariate fMRI studies, showing that familiarity is represented in a broad range of brain regions, including frontal (IFG, MPFC and anterior cingulate cortex, ATL), middle-central (TPJ, MTG/STS), and posterior regions (OFA, FFA and precuneus)(Castro-Laguardia et al., 2023; Natu & O’Toole, 2015; Visconti di Oleggio Castello et al., 2017). Together these results support the idea that familiarity information is represented both in the core and extended face networks (for a review see Kovács, 2020). This, in turn, suggests that the emerging familiarity representation is related to the representational transfer from perceptual processing towards person memory, a process attributed to several medial- and anterior-temporal regions and the hippocampus as well (Deen et al., 2024; Landi et al., 2021; Quian Quiroga et al., 2023). While the spatial resolution of the EEG does not allow us drawing such conclusions (and calls for future neuroimaging studies), the relative late emergence of familiarity information is in accordance with the results of previous ERP studies (Wiese et al., 2019a,2019b) as well as with the above interpretation.

Given the importance of visual similarity in EEG representation (de la Torre-Ortiz & Ruotsalo, 2024; Wardle et al, 2016), it is essential to control for it when measuring representations with MVPA. In recent years, DNNs became important for measuring perceptual similarity (e.g., Schroff et al., 2015, Shoham et al., 2023) and outperformed other computational models in explaining representations (Dobs et al., 2022; Shoham et al., 2023). After partialing out DNN-based similarity, familiarity representations appeared cca. 100 ms later. This suggests that earlier representations are modulated by visual similarity, while those after 400 ms are independent of that, a result consistent with prior results, showing familiarity modulation of ID representation from 400 ms onwards (Dalski et al., 2022; Dobs et al. 2019; Li et al., 2022). Altogether, our results support the idea that robust, identity-independent face familiarity representations emerge from around 400 ms post stimulus-onset, across the entire brain.

### Early face identity representations

First, we estimated the classification capacity of a machine learning algorithm from the EEG for same-sex face ID pairs, sharing visual face-semantic biography associations. For this we trained and tested LOPO classification of ID on participant data, having identical face-person knowledge associations. This analysis shows how the training on visual face-semantic biography association leads to ID specific representations.

ID representation started from around 100 ms post-stimulus onset in bilateral posterior cortical regions. This onset is similar to that of prior studies, using famous (Ambrus et al., 2019; Dobs et al., 2019; Kovács, 2023), media or personally familiarized IDs (Ambrus et al., 2021). Importantly, the length of the ID representation in our study (up to 800 ms) is similar to what was seen for highly familiar famous faces (e.g., Kovács, 2023). The spatial distribution of ID representation shows the strong involvement of posterior regions, in line with that of famous (Ambrus et al., 2019) as well as with media-familiarized faces (Ambrus et al., 2021). These results suggest that a brief familiarization training is sufficient to induce robust ID representations, comparable to those of previously familiarized famous faces or to those, resulting from longer media familiarization.

Results from the temporal generalization analysis suggest a different temporal dynamics of ID representations across scalp regions and imply that a cascade of processing from lower- to higher-level information integration is responsible for correct ID representation. The early, and temporally specific decoding in the left posterior ROI suggests transient, rapidly evolving ID representations. In contrast, these early transient ID representations were absent over the right hemisphere posterior ROIs, but the broadly generalized, stable ID representations emerged earlier there than over the left side. This hemispheric asymmetry is in line with prior studies suggesting functional dissociation between the two fusiform gyri, where the right FFA is involved in deeper cognitive processing of faces, while the left relates to more rapid, image-level face feature processing (Frässle et al., 2016; Meng et al., 2012).

The right central ROI showed a different temporal dynamic pattern compared to the right posterior region. This suggests different ID processing in these regions. We hypothesize that it may reflect the activation of cortical regions such as the STS and ATL, which have been suggested to play a role in the encoding of dynamic face features and person-related semantic knowledge respectively (Axelrod & Yovel, 2013; Collins & Olson, 2014; Duchaine & Yovel, 2015; Kovács, 2020; Lambon Ralph, 2014). The late and broadly generalized information over the left anterior ROI suggests stable higher-order representations, and the engagement of cortical regions associated with facial semantic and social information processing, including MPFC and IFG (Bobes et al., 2013; Frith & Frith, 1999; Cloutier et al., 2011).

Taken together, our results support the theory that ID processing occurs in a cascade across over several, temporally overlapping stages, from perceptual encoding over the left posterior cortex, to stable bilateral visual ID representations, towards higher-order semantic and conceptual processing in middle and anterior regions. This processing framework supports the hierarchical, distributed model of face ID representation involving both core and extended face-processing networks (for review, see Kovács, 2020).

### Person knowledge modulates visual identity representation

To estimate the role of person knowledge in ID representation, we performed LOPO classification for same-sex ID pairs, without shared biographical information as well. For this, we trained the algorithm on participant data having a given face-person knowledge association and tested it on that of those participants who had learned the associations in an opposite way (cross-group LOPO comparisons, see the Methods section and Figure 3b). We hypothesized that if the processing is sensitive to person knowledge, decoding accuracies will be reduced, when compared to the results of the within-group classification. Our results confirmed this hypothesis showing significantly lower classification performance between 250 – 370 ms over the right posterior electrode cluster. Moreover, the cross-group temporal generalization analysis revealed reduced decoding accuracy in the left anterior, right central and right posterior regions, as well as the reduction of off-diagonal generalization between 160 – 300 ms. These findings suggest that person-related biographical knowledge modulates strongly visual identity representations.

Previous univariate electrophysiological studies tested the neural correlates of ID-specific semantic and associative information processing. Our current results are at odds with these studies from many aspects. First, the first ERP component that shows clear access to person-specific semantic knowledge is later, corresponding to the N400 (e.g., Schweinberger & Burton, 2003; Wiese & Schweinberger, 2011, 2015; Wiese et al., 2024). Our results suggest that person-knowledge modulates representations at an earlier stage, beginning around 250 ms post-stimulus onset in the right posterior region. This time-window is the closest to the N250 ERP component, which peaks around 250 – 350 ms and is assumed to be generated by the neurons of the fusiform gyrus (Kaufmann et al., 2009). On the one hand, the N250 has been shown to exhibit enhanced responses to familiar faces (Schweinberger & Neumann, 2016; Wiese et al., 2024). On the other hand, the N250 is suggested to reflect the neural activity of image-invariant ID representations, given that the enhanced activation for familiar IDs persists across different exemplars of the same ID (Kaufmann et al., 2009). However, in contrast to our findings, previous ERP studies have failed to detect modulations of N250 amplitude by the associated semantic information (Kaufmann et al., 2009). This disagreement may be attributed to the inferior sensitivity of univariate methods when compared to MVPA. Indeed, information is encoded in a distributed activity pattern, rather than in the amplitude of evoked responses (Davidesco et al., 2014). Accordingly, prior MVPA studies have shown that ID representation starts around 200 ms (Ghuman et al., 2014; Vida et al., 2016) and suggested that person knowledge in the form of familiarity enhances ID representations within this time-range (Ambrus et al. 2021; Kovács et al. 2023). Notably, this time window is earlier than the emergence of ID-independent familiarity representations (Dalski et al., 2022; Dobs et al., 2019; Li et al., 2022), suggesting that the enhanced distinctiveness of ID representations is not related to the stronger familiarity representation, but better explained by the modulation of associated person knowledge, supporting the idea that such information modulates visual representations of familiar face identities and arguing for the separate mechanisms behind familiarity and ID representations further (Renoult et al., 2019; Rugg & Yonelinas, 2003; Kovács, 2020; Yonelinas, 2002).

The person knowledge modulation of ID representation was found in the right posterior electrode cluster in both decoding and temporal generalization analyses. The modulation aligns well with previous neuroimaging studies, which found that conceptual knowledge enhances the distinctiveness of visual face representations in corresponding regions (Brosch et al., 2013; Stolier & Freeman, 2016). Furthermore, fMRI MVPA studies have demonstrated that person-related information, such as personality traits, is encoded in posterior regions, specifically in the FFA (Cao et al., 2020; Tsantani et al., 2021). Given the well-established role of posterior regions, such as the FFA, in visual ID processing (Kanwisher & Yovel, 2006; Rotshtein et al, 2005), these findings, together with the cortical distribution of the observed modulation effect in our study, support the view that person-related knowledge modulates face ID representations already at this early processing stage. This conclusion aligns well with the dynamic model of face perception (Freeman & Ambady, 2011), suggesting that higher-order, social-cognitive processes also impact lower-level visual processes as well (Stolier & Freeman, 2016).

However, a previous fMRI study suggested that the association of detailed biographical information leads to higher representation similarity when compared to simple, face-name associations (Verosky et al., 2013). This is at odds with our results, showing enhanced decoding for faces, associated to person knowledge. The apparent contradiction may be due to the fact that the enhanced representation was only found in a limited time-window in our study, which might be overlooked by fMRI. Further, it is possible that prior knowledge associations do not enhance neural dissimilarity between face representations universally and the effect depends on the similarity of the associated conceptual content (Bein et al., 2020). When the associated conceptual knowledge is unique, such as social bias and stereotypes (e.g., Brosch et al., 2013) or biographical knowledge, conceptual differences may shape the neural representations of visual identity by increasing their distinctiveness.

Finally, the temporal generalization analysis revealed that person-knowledge modulates ID representations in the left anterior and right central regions. The reduction of temporal generalization accuracy in cross association group comparisons may reflect the modulating effect of the reversed person-knowledge associations in the cross-group analysis. Intriguingly, ID representation remained unaffected by the reversed person-related knowledge associations in the left posterior region. These findings support the idea of hemispherical functional asymmetry in the face-processing network, especially in occipito-temporal regions (Bowers et al., 1985; Frässle et al., 2016; Meng et al., 2012). In line with prior studies which suggest that the right FFA encodes face identity (Ma & Han, 2012) and personality trait representations (Cao et al., 2020). Importantly we show that these right posterior ID representations are sensitive to person-related knowledge as well, while those of the left hemisphere remain unaffected. Our findings call for future EEG and fMRI studies to clarify the functional roles and connectivity of bilateral posterior regions, especially FFA, during the progress of integrating perceptual and conceptual information in face familiarization and recognition (Kovács, 2020).

### Conclusion

In conclusion, the current study reveals how person-related conceptual and biographical knowledge modulates the spatiotemporal dynamics of familiar ID representations. Using multivariate EEG approach, we found that person knowledge modulates face ID processing between 250 – 370 ms post-stimulus onset, over the right posterior cortical electrodes. This suggests the top-down, person recognition memory related modulation of visual identity processing. Our findings support current dynamic face perception models and help uncovering the mechanism of how semantic and biographical information shape identity representations.

## Funding and Acknowledgements

Y.S is supported by the China Scholarship Council (CSC) scholarship (202308330055).

