## Supplementary Materials for "It matters who you are: Biography modulates the neural dynamics of facial identity representation"

| Nationality | Gender | Text |
| --- | --- | --- |
| German | Male | Karl Müller, 34 years old, is a pragmatic and disciplined middle school teacher from Frankfurt, Germany. Known for his punctuality, strictness and bad sense of humor. He enjoys hiking in the Black Forest and playing football with friends |
| German | Female | Anna Schneider, 31 years old, is an organized and thoughtful engineer from Bonn, Germany. She is known for her carefulness, perfectionism and hardworking. She loves cycling along the river, and gardening in her yard. |
| French | Male | Jean Laurent, 36 years old, is a sociable chef from Lyon, France. He is known for his charm, elegant and passionate. He likes playing guitar, reading books and cooking traditional French cuisines. |
| French | Female | Louise Cosette, 29 years old, is a creative and outgoing designer from Paris, France. She is known for her warm and active personality. She enjoys art and fashion and spending her weekends watching classical movies and visiting museums. |

Appendix S1. full text of all four biographical information paragraphs

| Number | Question |
| --- | --- |
| 1 | What is his/her name? |
| 2 | Where does he/she come from? |
| 3 | What is his/her job? |
| 4  5 | What is his/her hobby?  What is his/her personality? |

Appendix S2. full text of all four biographical information paragraphs

| ROI | Onset (ms) | Peak (ms) | End (ms) | Cluster-based *p*-value | |
| --- | --- | --- | --- | --- | --- |
| All electrodes | 300 | 550 | 1200 | | < 0.001 |
| Left anterior | 390 | 1050 | 1200 | | < 0.001 |
| Right anterior | 440 | 810 | 1200 | | < 0.001 |
| Left central | 390 | 540; 960 | 1200 | | < 0.001; < 0.001 |
| Right central | 440 | 560 | 1200 | | < 0.001 |
| Left posterior | 250 | 550; 870 | 1200 | | < 0.001; < 0.001 |
| Right posterior | 440 | 550; 1080 | 1200 | | < 0.001; < 0.001 |

Supplementary Table 1. Results of the time-resolved RSA analysis of familiarity information, without partialling out DNN-based similarity. Clusters marked as significant on Fig. 4(yellow curve). Two-tailed one-sample cluster-based permutation test against zero correlation.

| ROI | Onset (ms) | Peak (ms) | End (ms) | Cluster-based *p*-value |
| --- | --- | --- | --- | --- |
| All electrodes | 440 | 560 | 1200 | < 0.001 |
| Left anterior | 400 | 1050 | 1200 | < 0.001 |
| Right anterior | 450 | 1020 | 1200 | < 0.001 |
| Left central | 480 | 610; 960; 1170 | 1200 | 0.004; 0.007; 0.009 |
| Right central | 460 | 570; 1150 | 1200 | 0.001; 0.001 |
| Left posterior | 370 | 550; 870 | 1200 | < 0.001; < 0.001 |
| Right posterior | 450 | 550; 1040 | 1130 | < 0.001; < 0.001 |

Supplementary Table 2. Results of the time-resolved RSA analysis of familiarity information after partialling out DNN-based similarity. Clusters marked as significant on Fig. 4(blue curve). Two-tailed one-sample cluster-based permutation test against zero correlation.
